# Microbiome-to-Nigrostriatal Vulnerability Mapping Prioritizes SCFA-Linked Gut-Brain Axes in Parkinson’s Disease

**DOI:** 10.64898/2026.07.11.737943

**Authors:** Nabanita Ghosh

**Affiliations:** Department of Zoology, Maulana Azad College, Kolkata, West Bengal, India

**Author notes:** **Correspondence** Correspondence and requests for materials should be addressed to Nabanita Ghosh.

**Keywords:** Parkinson’s disease, gut microbiome, short-chain fatty acids, butyrate, IL1B, gut-brain axis, transcriptomics, systems biology, evidence integration

## Abstract

**Background:** Parkinson’s disease (PD) is increasingly recognized as a multisystem neurodegenerative disorder in which gastrointestinal dysfunction, microbial ecology, immune signaling, and nigrostriatal vulnerability intersect. Microbiome studies have identified PD-associated gut microbial alterations, but translating these observations into host-relevant biological axes requires integration across microbial functions, metabolites, host genes, and brain-region transcriptomic context.

**Methods:** MiNi-PD was developed as a lightweight, fully in silico, table-level prioritization workflow integrating processed PD microbiome features, curated microbe–metabolite–host-gene associations from gutMGene v2.0, and processed substantia nigra and putamen transcriptomic support from GSE136666. The framework ranks microbe–metabolite–host-gene axes using exact-curated metabolite-mediated evidence, brain-region support, and a normalized Nigrostriatal Gut-Brain Convergence Score (NGBCS).

**Results:** Strict evidence filtering retained 639 primary exact-curated metabolite-mediated axes. Of these, 12 had FDR-level brain transcriptomic support and 65 had nominal-or-stronger support. IL1B emerged as the principal FDR-level host-gene convergence point. SCFA-linked metabolites dominated the primary exact-curated set, including 387 butyrate axes and 5 3-indolepropionic-acid axes. The FDR-supported axes reached the maximum NGBCS value of 5.0, while the top nominally supported axes reached the high-priority NGBCS value of 4.5.

**Conclusion:** MiNi-PD identifies a focused SCFA-linked gut-brain signature in PD, highlighting butyrate- and 3-indolepropionic-acid-associated links to IL1B as a biologically meaningful substantia nigra-supported inflammatory axis. The NGBCS framework provides an accessible and interpretable route for prioritizing metabolite-mediated gut-brain mechanisms for downstream validation.

## 1 Introduction

Parkinson’s disease (PD) is a progressive neurodegenerative disorder classically defined by bradykinesia, rigidity, resting tremor, and postural instability, but the clinical syndrome extends beyond motor impairment. Non-motor features, including autonomic dysfunction, sleep disturbance, neuropsychiatric symptoms, olfactory dysfunction, and gastrointestinal manifestations, contribute substantially to disease burden and may emerge before a motor diagnosis is established. Neuropathological staging frameworks and gut-first hypotheses have therefore motivated sustained interest in peripheral and enteric contributors to PD vulnerability [4, 5].

Gastrointestinal symptoms, particularly constipation, are common in PD and can precede motor manifestations by years. Their early occurrence places gut biology within the temporal and biological landscape of PD progression and supports the relevance of gut-brain communication as a disease-contextual axis. Experimental and observational studies have suggested that microbial communities, gut barrier biology, immune signaling, and metabolite exposure may influence host inflammatory states and neuronal vulnerability, making the gut an important source of mechanisms for prioritization [5, 6].

Human microbiome studies have reported altered gut microbial composition in PD and related prodromal states [6, 7, 1]. However, taxonomic findings alone are difficult to translate into host-relevant hypotheses. A microbial species can participate in multiple metabolic activities, and the biological implication of a taxonomic association depends on metabolites, host target genes, tissue context, and disease-relevant downstream pathways. Therefore, a systems-level framework is needed to connect microbial features to potential host response axes.

Microbial metabolites offer a plausible bridge between gut microbial variation and host biology. Short-chain fatty acids (SCFAs), including butyrate, acetate, and propionate, can influence epithelial, immune, and epigenetic responses. Tryptophan and indole derivatives, succinate, immune ligands, and inflammatory mediators provide additional routes through which microbial communities may be linked to host transcriptional programs. Prior work has reported altered fecal SCFA profiles in PD [8], but metabolite-centered prioritization still requires integration with curated host-gene evidence and PD-relevant brain transcriptomic support.

MiNi-PD was designed to address this gap using a reproducible and computationally light table-level workflow. MiNi-PD links processed PD microbiome result tables to curated microbe–metabolite–host-gene relationships and then evaluates whether host genes show transcriptomic support in substantia nigra or putamen. This design supports interpretable evidence integration and produces ranked microbe–metabolite–host-gene axes for biological prioritization.

The objective of the present study is to describe MiNi-PD and its strict evidence-tiered output for PD gut-brain axis prioritization. We tested whether PD-associated microbial features converge on curated metabolite-mediated host-gene axes with brain-region transcriptomic support using the Wallen et al. PD microbiome resource, gutMGene v2.0, and GSE136666 brain-region transcriptomic data. The analysis identifies an SCFA-dominated signal and prioritizes butyrate- and 3-indolepropionic-acid-associated links to IL1B as a focused inflammatory axis with substantia nigra support.

## 2 Methods

### 2.1 Study design

MiNi-PD is a fully in silico, table-level prioritization workflow that integrates processed public microbiome, curated microbe–metabolite–host-gene, and processed brain transcriptomic tables. The workflow ranks metabolite-mediated gut-brain axes using transparent evidence components derived from public source tables and a normalized NGBCS index.

### 2.2 Data sources

The primary microbiome evidence layer used Wallen et al. 2022 Nature Communications PD metagenomics supplementary result tables [1]. The species layer used Supplementary Data 1, the microbial gene/KO layer used Supplementary Data 9, and the MetaCyc/pathway layer used Supplementary Data 10. The microbe–metabolite–host-gene bridge used gutMGene v2.0 component CSV files [2]: gutMGene1.csv for microbe–metabolite relationships, gutMGene2.csv for metabolite–host-gene relationships, and gutMGene3.csv for direct microbe–host-gene relationships. Brain transcriptomic support used processed GSE136666 / Xicoy et al. tables [3]: Spreadsheet S2 RNAseq substantia nigra and Spreadsheet S4 RNAseq putamen.

### 2.3 Data conversion and provenance

Downloaded Excel and CSV files were converted to tab-separated values (TSV) for stable local processing. The source manifest and curation log were maintained in the project metadata directory. Conversion included format harmonization, column-name whitespace stripping, and removal of fully empty rows or columns.

### 2.4 gutMGene harmonization

The gutMGene component files were harmonized using the relationships represented in the source tables. The microbe–metabolite component contained 741 rows, the metabolite–host-gene component contained 244 rows, and the direct microbe–host-gene component contained 215 rows. Metabolite-mediated joins were performed through exact normalized metabolite names. There were 45 unique normalized metabolites with host-gene evidence, producing 3069 joined rows before deduplication. The final harmonized output contained 3284 rows after deduplication. Direct microbe–host-gene rows were retained with a blank metabolite field as a supplementary evidence tier, while explicit microbe–metabolite–host-gene axes formed the primary metabolite-mediated evidence tier.

### 2.5 Microbial feature standardization

Wallen species, gene/KO, and pathway tables were standardized into a common microbial-feature format with harmonized feature identifiers, feature names, feature type, direction in PD, effect size, p value, FDR/q value, source table, and threshold status where available. The Wallen species table contained 720 rows, the Wallen genes/KO table contained 8529 rows, and the Wallen pathways table contained 512 rows. The standardized microbial feature table contained 9761 rows.

### 2.6 Microbial module annotation

Microbial features were assigned to biological modules using transparent rule-based annotation. SCFA annotations included metabolites and taxa linked to butyrate, acetate, propionate, and related short-chain fatty-acid biology. Oxidative-stress and inflammatory annotations used biological pathway terms, including oxidative stress, reactive oxygen, peroxide, peroxidase, superoxide, glutaredoxin, thioredoxin, redox, oxidoreductase, and inflammatory signaling terms. SCFA taxa and metabolites, including butyrate-related taxa such as *Roseburia, Anaerostipes, Faecalibacterium, Eubacterium, Butyricimonas*, and *Butyrivibrio*, were assigned to the SCFA module when supported by the annotation rules.

### 2.7 Brain transcriptomic support

Processed GSE136666 substantia nigra and putamen tables were used to assess brain transcriptomic support for gut-derived host genes. Each regional table contained 19169 rows. Host-gene support was categorized as FDR-level support when the adjusted significance value passed the configured threshold and as nominal support when nominal significance was present without FDR-level support. This tiered evidence structure allowed FDR-level and nominal transcriptomic support to be reported separately.

### 2.8 NGBCS scoring and strict evidence tiers

The Nigrostriatal Gut-Brain Convergence Score (NGBCS) is a normalized evidence-convergence index developed to rank microbe–metabolite–host-gene axes by the amount and specificity of support connecting PD-associated microbiome features to nigrostriatal transcriptomic context. The index is bounded from 0 to 5, where 0 indicates no retained evidence for a complete gut-brain axis and 5 represents the maximum convergence tier in this implementation. NGBCS summarizes microbial threshold support, interpretable microbial direction, exact-curated microbial matching, explicit metabolite mediation, host-gene evidence, brain transcriptomic support, pathway relevance, and nigrostriatal specificity. For interpretation, axes with NGBCS *≥* 4.5 were considered high-priority evidence-supported axes in the main analysis, while axes with NGBCS below 4.5 were retained as lower-priority or background results and were not emphasized in the main evidence tables. Within the high-priority range, NGBCS = 5.0 denotes the strongest evidence tier, corresponding to exact-curated metabolite-mediated axes with FDR-level brain transcriptomic support. NGBCS = 4.5 denotes a strong but secondary tier, corresponding to exact-curated metabolite-mediated axes with nominal brain transcriptomic support. Thus, the uniform NGBCS values shown in Table 1 and Table 2 reflect their evidence tiers: Table 1 contains maximum-score axes and Table 2 contains high-priority nominally supported axes. Exact-curated axes were treated as the primary evidence tier. Genus-level curated axes were retained as an exploratory tier. Direct microbe–host-gene evidence was retained as a supplementary tier. Primary interpretation prioritized exact-curated metabolite-mediated axes.

**Table 1:** FDR-supported exact microbe–metabolite–host-gene axes identified by MiNi-PD. NGBCS ranges from 0 to 5; all rows reach the maximum score of 5.0, representing the strongest evidence-convergence tier.

| Microbe | Metabolite | Host gene | Direction | Module | NGBCS | Brain regions | Interpretation note |
| --- | --- | --- | --- | --- | --- | --- | --- |
| *Anaerostipes hadrus* | Butyrate | IL1B | down | SCFA | 5 | putamen;<br>substantia_nigra | Candidate exact-curated metabolite-mediated axis with brain transcriptomic support. |
| *Eubacterium limosum* | Butyrate | IL1B | up | SCFA | 5 | putamen;<br>substantia_nigra | Candidate exact-curated metabolite-mediated axis with brain transcriptomic support. |
| *Eubacterium ramulus* | Butyrate | IL1B | down | SCFA | 5 | putamen;<br>substantia_nigra | Candidate exact-curated metabolite-mediated axis with brain transcriptomic support. |
| *Faecalibacterium prausnitzii* | 3-Indolepropionic acid | IL1B | down | SCFA | 5 | putamen;<br>substantia_nigra | Candidate exact-curated metabolite-mediated axis with brain transcriptomic support. |
| *Faecalibacterium prausnitzii* | Butyrate | IL1B | down | SCFA | 5 | putamen;<br>substantia_nigra | Candidate exact-curated metabolite-mediated axis with brain transcriptomic support. |
| *Roseburia intestinalis* | Butyrate | IL1B | down | SCFA | 5 | putamen;<br>substantia_nigra | Candidate exact-curated metabolite-mediated axis with brain transcriptomic support. |
| *Roseburia inulinivorans* | Butyrate | IL1B | down | SCFA | 5 | putamen;<br>substantia_nigra | Candidate exact-curated metabolite-mediated axis with brain transcriptomic support. |
| *Alistipes indistinctus* | Succinate | IL1B | up | Unassigned | 5 | putamen;<br>substantia_nigra | Candidate exact-curated metabolite-mediated axis with brain transcriptomic support. |
| *Bifidobacterium longum* | Butyrate | IL1B | up | Unassigned | 5 | putamen;<br>substantia_nigra | Candidate exact-curated metabolite-mediated axis with brain transcriptomic support. |
| *Christensenella minuta* | Butyrate | IL1B | up | Unassigned | 5 | putamen;<br>substantia_nigra | Candidate exact-curated metabolite-mediated axis with brain transcriptomic support. |
| *Parabacteroides distasonis* | Succinate | IL1B | up | Unassigned | 5 | putamen;<br>substantia_nigra | Candidate exact-curated metabolite-mediated axis with brain transcriptomic support. |
| *Streptococcus vestibularis* | Butyrate | IL1B | up | Unassigned | 5 | putamen;<br>substantia_nigra | Candidate exact-curated metabolite-mediated axis with brain transcriptomic support. |

**Table 2:** Top nominally brain-supported exact metabolite-mediated axes. NGBCS ranges from 0 to 5; rows shown here reach 4.5, the high-priority nominalsupport tier.

| Microbe | Metabolite | Host gene | Direction | Module | NGBCS | Brain regions |
| --- | --- | --- | --- | --- | --- | --- |
| *Anaerostipes hadrus* | Butyrate | CHGA | down | SCFA | 4.5 | putamen |
| *Eubacterium limosum* | Butyrate | CHGA | up | SCFA | 4.5 | putamen |
| *Eubacterium ramulus* | Butyrate | CHGA | down | SCFA | 4.5 | putamen |
| *Faecalibacterium prausnitzii* | Butyrate | CHGA | down | SCFA | 4.5 | putamen |
| *Roseburia intestinalis* | Butyrate | CHGA | down | SCFA | 4.5 | putamen |
| *Roseburia inulinivorans* | Butyrate | CHGA | down | SCFA | 4.5 | putamen |
| *Anaerostipes hadrus* | Butyrate | GCK | down | SCFA | 4.5 | putamen |
| *Eubacterium limosum* | Butyrate | GCK | up | SCFA | 4.5 | putamen |
| *Eubacterium ramulus* | Butyrate | GCK | down | SCFA | 4.5 | putamen |
| *Faecalibacterium prausnitzii* | Butyrate | GCK | down | SCFA | 4.5 | putamen |
| *Roseburia intestinalis* | Butyrate | GCK | down | SCFA | 4.5 | putamen |
| *Roseburia inulinivorans* | Butyrate | GCK | down | SCFA | 4.5 | putamen |
| *Anaerostipes hadrus* | Butyrate | HDAC1 | down | SCFA | 4.5 | putamen |
| *Eubacterium limosum* | Butyrate | HDAC1 | up | SCFA | 4.5 | putamen |
| *Eubacterium ramulus* | Butyrate | HDAC1 | down | SCFA | 4.5 | putamen |
| *Faecalibacterium prausnitzii* | Butyrate | HDAC1 | down | SCFA | 4.5 | putamen |
| *Roseburia intestinalis* | Butyrate | HDAC1 | down | SCFA | 4.5 | putamen |
| *Roseburia inulinivorans* | Butyrate | HDAC1 | down | SCFA | 4.5 | putamen |
| *Faecalibacterium prausnitzii* | 3-Indolepropionic acid | IL6 | down | SCFA | 4.5 | substantia_nigra |
| *Anaerostipes hadrus* | Acetate | IL6 | down | SCFA | 4.5 | substantia_nigra |

Nominal support is reported separately from FDR-level support; NGBCS = 4.5 denotes the high-priority nominal-support tier.

### 2.9 Strict evidence filtering

Strict evidence filtering was applied after NGBCS scoring. The full NGBCS output contained 31,746 total axes. Strict exact-curated filtering retained 679 axes. Requiring explicit metabolite mediation produced 639 primary exact metabolite-mediated axes. Among these, 12 had FDR-level brain transcriptomic support, 65 had nominal-or-stronger brain transcriptomic support, and 574 had no detected brain transcriptomic support under the current criteria. Genus-level and direct microbe–host-gene rows were kept separate as supplementary evidence tiers.

### 2.10 Statistical framework

FDR-level terminology follows the Benjamini-Hochberg false-discovery-rate framework [9]. Brain transcriptomic support was summarized at host-gene and module levels. FDR-level and nominal evidence were retained as distinct support tiers, and NGBCS *≥* 4.5 was used as the operational high-priority score range for the main evidence tables.

## 3 Results

### 3.1 MiNi-PD prioritizes exact-curated metabolite-mediated gut-brain axes

MiNi-PD linked processed PD microbiome results [1], gutMGene v2.0 microbe–metabolite–host-gene associations [2], and processed GSE136666 substantia nigra and putamen transcriptomic tables [3]. After strict evidence filtering, the primary evidence set contained 639 exact-curated metabolite-mediated axes. These rows required exact microbial matching, explicit metabolite mediation, and Wallen-threshold microbial evidence. Genus-level matches and direct microbe–host-gene rows were retained only as supplementary exploratory evidence.

### 3.2 Brain-supported axes identify IL1B as a principal convergence point

Among the 639 primary exact-curated axes, 12 had FDR-level brain transcriptomic support and 65 had nominal-or-stronger brain transcriptomic support. Table 1 summarizes the FDR-supported exact axes, and Table 2 summarizes the nominally supported exact axes. IL1B was the strongest host-gene convergence point and the only gene with FDR-level substantia nigra transcriptomic support; host-gene support is summarized in Table 3. The axes summarized in Table 1 reached the maximum NGBCS value of 5.0, indicating the strongest evidence-convergence tier. The nominally supported axes summarized in Table 2 reached NGBCS values of 4.5, meeting the high-priority score threshold while remaining distinct from the FDR-supported tier.

**Table 3:** Host-gene support summary for primary exact metabolite-mediated axes.

| Host gene | Exact axes | Microbes | Metabolites | FDR | Nominal | Brain regions | Suggested role |
| --- | --- | --- | --- | --- | --- | --- | --- |
| IL1B | 12 | 11 | 3 | Yes | Yes | putamen;<br>substantia_nigra | inflammatory cytokine / innate immune activation |
| IL6 | 20 | 10 | 4 | No | Yes | substantia_nigra | inflammatory cytokine |
| CHGA | 9 | 9 | 1 | No | Yes | putamen | neuroendocrine / secretory pathway |
| GCK | 9 | 9 | 1 | No | Yes | putamen | metabolic regulation |
| HDAC1 | 9 | 9 | 1 | No | Yes | putamen | epigenetic regulation / SCFA-related host response |
| MAPK3 | 2 | 2 | 1 | No | Yes | putamen;<br>substantia_nigra | MAPK signaling |
| TLR9 | 2 | 2 | 1 | No | Yes | substantia_nigra | innate immune receptor |
| NFKB1 | 1 | 1 | 1 | No | Yes | putamen | NF-κB signaling |
| RHEB | 1 | 1 | 1 | No | Yes | substantia_nigra | mTOR/autophagy-related signaling |

### 3.3 SCFA-linked metabolites dominate the primary signal

SCFA-associated metabolites dominated the primary exact-curated axis set. Table 4 summarizes metabolite-level support using compact counts; detailed metabolite-linked microbe and host-gene lists are provided in Supplementary Table S3. Butyrate accounted for 387 axes, acetate for 168 axes, propionate for 38 axes, succinate for 16 axes, and 3-indolepropionic acid for 5 axes. Table 5 summarizes microbe-level support using compact counts; detailed microbe-linked metabolite and host-gene lists are provided in Supplementary Table S4. The highest-count microbial features included *Eubacterium limosum, Eubacterium ramulus, Bifidobacterium longum, Faecalibacterium prausnitzii, Anaerostipes hadrus, Christensenella minuta, Roseburia intestinalis*, and *Roseburia inulinivorans*.

**Table 4:** Compact metabolite-level support summary.

| Metabolite | Exact axes | Microbes, n | Host genes, n | FDR genes | Nominal genes | Module |
| --- | --- | --- | --- | --- | --- | --- |
| Butyrate | 387 | 9 | 43 | IL1B | CHGA;<br>GCK;<br>HDAC1;<br>IL1B;<br>IL6 | SCFA |
| Acetate | 168 | 8 | 21 |  | IL6 | SCFA |
| Propionate | 38 | 2 | 19 |  | IL6;<br>TLR9 | SCFA |
| Succinate | 16 | 2 | 8 | IL1B | IL1B;<br>MAPK3 | Unassigned |
| 3-Indolepropionic acid | 5 | 1 | 5 | IL1B | IL1B;<br>IL6;<br>NFKB1 | SCFA |
| Caffeic Acid | 8 | 2 | 4 |  |  | Unassigned |
| 3-(4-Hydroxyphenyl)propionic acid | 4 | 1 | 4 |  |  | SCFA |
| 4-Hydroxyphenylacetic acid | 4 | 1 | 4 |  |  | SCFA |
| Genipin | 4 | 1 | 4 |  |  | Unassigned |
| Hydroquinone | 3 | 1 | 3 |  |  | Unassigned |
| 5-hydroxy-1H-imidazole-4-carboxamide | 2 | 1 | 2 |  | RHEB | Unassigned |

Detailed microbe and host-gene lists are provided in Supplementary Table S3.

**Table 5:** Compact microbe-level support summary.

| Microbe | Direction | Module | Exact axes | Metabolites, n | Host genes, n | FDR genes | Nominal genes |
| --- | --- | --- | --- | --- | --- | --- | --- |
| *Eubacterium limosum* | up | SCFA | 83 | 3 | 53 | IL1B | CHGA;<br>GCK;<br>HDAC1;<br>IL1B;<br>IL6;<br>TLR9 |
| *Eubacterium ramulus* | down | SCFA | 72 | 4 | 51 | IL1B | CHGA;<br>GCK;<br>HDAC1;<br>IL1B;<br>IL6 |
| *Bifidobacterium longum* | up | Unassigned | 71 | 4 | 52 | IL1B | CHGA;<br>GCK;<br>HDAC1;<br>IL1B;<br>IL6 |
| *Faecalibacterium prausnitzii* | down | SCFA | 69 | 3 | 49 | IL1B | CHGA;<br>GCK;<br>HDAC1;<br>IL1B;<br>IL6;<br>NFKB1 |
| *Anaerostipes hadrus* | down | SCFA | 64 | 2 | 48 | IL1B | CHGA;<br>GCK;<br>HDAC1;<br>IL1B;<br>IL6 |
| *Christensenella minuta* | up | Unassigned | 64 | 2 | 48 | IL1B | CHGA;<br>GCK;<br>HDAC1;<br>IL1B;<br>IL6 |
| *Roseburia intestinalis* | down | SCFA | 64 | 2 | 48 | IL1B | CHGA;<br>GCK;<br>HDAC1;<br>IL1B;<br>IL6 |
| *Roseburia inulinivorans* | down | SCFA | 62 | 2 | 48 | IL1B | CHGA;<br>GCK;<br>HDAC1;<br>IL1B;<br>IL6;<br>TLR9 |
| *Streptococcus vestibularis* | up | Unassigned | 43 | 1 | 43 | IL1B | CHGA;<br>GCK;<br>HDAC1;<br>IL1B;<br>IL6 |
| *Streptococcus australis* | down | Unassigned | 21 | 1 | 21 |  | IL6 |
| *Alistipes indistinctus* | up | Unassigned | 8 | 1 | 8 | IL1B | IL1B;<br>MAPK3 |
| *Parabacteroides distasonis* | up | Unassigned | 8 | 1 | 8 | IL1B | IL1B;<br>MAPK3 |
| *Escherichia coli* | up | Unassigned | 4 | 1 | 4 |  |  |
| *Lactobacillus gasseri* | up | Unassigned | 4 | 1 | 4 |  |  |
| *Streptococcus mutans* | up | Unassigned | 2 | 1 | 2 |  | RHEB |

Detailed metabolite and host-gene lists are provided in Supplementary Table S4.

### 3.4 Nominal host-gene support provides biological context

In addition to the FDR-level IL1B signal, nominal transcriptomic support was observed for IL6, CHGA, GCK, HDAC1, MAPK3, TLR9, and NFKB1. These genes provide cautious biological context spanning inflammatory cytokine signaling, NF-*κ*B-related signaling, epigenetic regulation, metabolic regulation, MAPK signaling, neuroendocrine/secretory biology, and innate immune receptor signaling. These nominally supported genes add biological context to the IL1B-centered FDR-level signal and expand the inflammatory-metabolic profile of the prioritized SCFA-linked axes.

### 3.5 Exploratory evidence is retained separately

Genus-level axes and direct microbe–host-gene rows were not included in the primary evidence tier. Genus-level rows are less specific than exact-curated microbial matches, and direct microbe–hostgene rows do not provide explicit metabolite mediation. Both evidence classes are therefore retained as supplementary material to provide broader biological context while keeping the main evidence tier specific to exact-curated metabolite-mediated axes.

### 3.6 Integrated result

Together, these results define a focused SCFA-linked gut-brain signature in PD. MiNi-PD prioritized butyrate- and 3-indolepropionic-acid-associated links to IL1B as the central inflammatory axis, supported by exact-curated metabolite mediation, high NGBCS values, and substantia nigra transcriptomic evidence.

## 4 Discussion

MiNi-PD identifies a focused SCFA-linked gut-brain signature in Parkinson’s disease by integrating PD microbiome result tables, curated microbe–metabolite–host-gene associations, and brain-region transcriptomic support. The principal finding is an IL1B-centered inflammatory convergence within exact-curated metabolite-mediated axes, supported at the FDR level in substantia nigra and represented by the maximum NGBCS tier. The explicit NGBCS range strengthens the interpretability of this result: scores span 0 to 5, axes with NGBCS *≥* 4.5 represent the high-priority evidence-supported range, FDR-supported axes reach the maximum score of 5.0, and nominally brain-supported axes reach the secondary high-priority score of 4.5. Therefore, the score values shown in Tables 1 and 2 are not arbitrary repeated numbers; they identify two biologically meaningful evidence tiers, separating the highest-confidence IL1B-centered FDR-supported axes from the broader nominally supported inflammatory and metabolic context. The dominance of butyrate-linked axes is an important feature of the result. Butyrate accounted for 387 primary exact-curated axes, while acetate, propionate, succinate, and 3-indolepropionic acid contributed additional metabolite-mediated links. These metabolites are biologically relevant because SCFAs and indole-derived metabolites are positioned at the interface of microbial ecology, epithelial regulation, immune signaling, and host transcriptional control; altered SCFA profiles and PD-associated gut microbial differences have been reported previously [8, 6, 1]. The prioritized microbial features, including *Faecalibacterium prausnitzii, Anaerostipes hadrus, Roseburia intestinalis, Roseburia inulinivorans, Eubacterium limosum*, and *Eubacterium ramulus*, provide a biologically plausible microbial context for the SCFA-centered signal. The IL1B result gives the prioritization a strong inflammatory anchor. IL1B is a central cytokine in innate immune activation, and experimental PD models have linked gut microbiota to neuroinflammatory states [5]. Its FDR-level substantia nigra support in GSE136666 distinguishes it from the broader set of nominally supported genes and makes it the most robust host-gene point in the present analysis [3]. The additional nominally supported genes, including IL6, CHGA, GCK, HDAC1, MAPK3, TLR9, and NFKB1, add a coherent secondary context spanning cytokine biology, NF-*κ*B-related signaling, MAPK signaling, epigenetic regulation, metabolic regulation, innate immune recognition, and neuroendocrine or secretory function. The value of MiNi-PD lies not only in the individual IL1B-centered result but also in the structure of the workflow: by separating exact-curated metabolite-mediated axes from genus-level and direct microbe–host-gene evidence, the analysis preserves a stringent primary result while retaining broader supplementary context. This organization is useful for PD gut-brain research because microbiome studies often produce large lists of taxa or pathways that are difficult to translate into host-relevant biology [6, 7, 1]. MiNi-PD condenses those inputs into interpretable microbial metabolite axes linked to brain-region gene support and ranks them through an explicit convergence score. The present findings also provide a practical framework for future biological testing. The most direct next step is targeted investigation of SCFA-linked axes involving butyrate, 3-indolepropionic acid, succinate, and IL1B-related inflammatory responses in independent PD microbiome cohorts, targeted metabolomics datasets, intestinal or immune-cell models, and brain-region or cell-type-aware transcriptomic resources. Overall, MiNi-PD strengthens the biological interpretation of PD gut microbiome data by connecting microbial features to metabolite-mediated host-gene axes and substantia nigra transcriptomic support across the three public evidence layers used here [1, 2, 3]. The study highlights an SCFA-linked, IL1B-centered inflammatory axis as a significant and actionable result for subsequent experimental and computational validation. The scope of the work is an integrative table-level prioritization analysis based on processed public microbiome, gutMGene, and brain transcriptomic resources. Its principal limitations are the absence of causal inference, direct metabolomics, experimental testing, raw-read reprocessing, and de novo cell-type analysis, together with possible gutMGene literature bias and the observation that module-level brain enrichment was not FDR-significant.

## 5 Ethics Statement

This study used publicly available processed or supplementary tables and did not involve new recruitment, intervention, raw human sequencing data generation, or access to identifiable participant-level records. No new ethics approval was required for this table-level secondary analysis.

## 6 Data Availability

All input data analyzed in this study were derived from publicly available processed or supplementary tables. The microbiome evidence layer used Wallen et al. 2022 supplementary species, KO/gene, and MetaCyc/pathway result tables. Microbe–metabolite–host-gene relationships were derived from gutMGene v2.0 component exports. Brain transcriptomic support was derived from processed GSE136666 / Xicoy et al. substantia nigra and putamen tables. Locally converted TSV inputs, source provenance notes, and curation records are documented in the MiNi-PD project metadata, hosted in Zenodo (https://doi.org/10.5281/zenodo.20192507).

## 7 Code Availability

The code is available in Zenodo repository (https://doi.org/10.5281/zenodo.20192507).

## 9 Author Contributions

Nabanita Ghosh conceived the study, designed the MiNi-PD workflow, executed the computational analysis, interpreted the results, and prepared the manuscript.

## 12 Tables

Main result tables are inserted below. Full TSV versions of Tables 1–5 are also provided in the accompanying tables directory. Large supplementary evidence-tier tables remain separate and are described in the Supplementary Material Description section.

## 13 Figures

**Figure 1.**
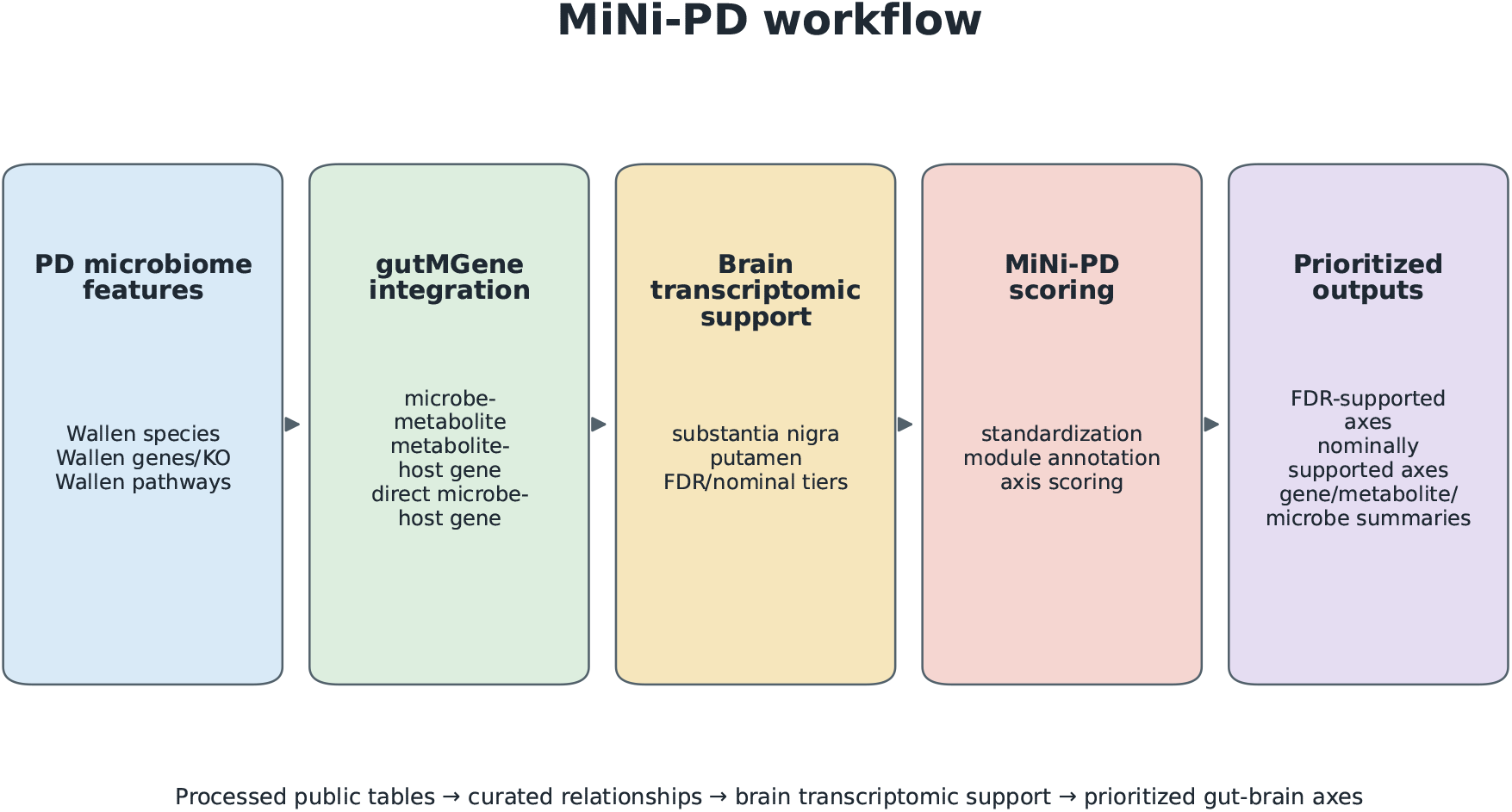
MiNi-PD workflow. MiNi-PD links processed PD microbiome tables, curated gutMGene microbe–metabolite–host-gene relationships, and GSE136666 brain transcriptomic tables to prioritize metabolite-mediated gut–brain axes.

**Figure 2.**
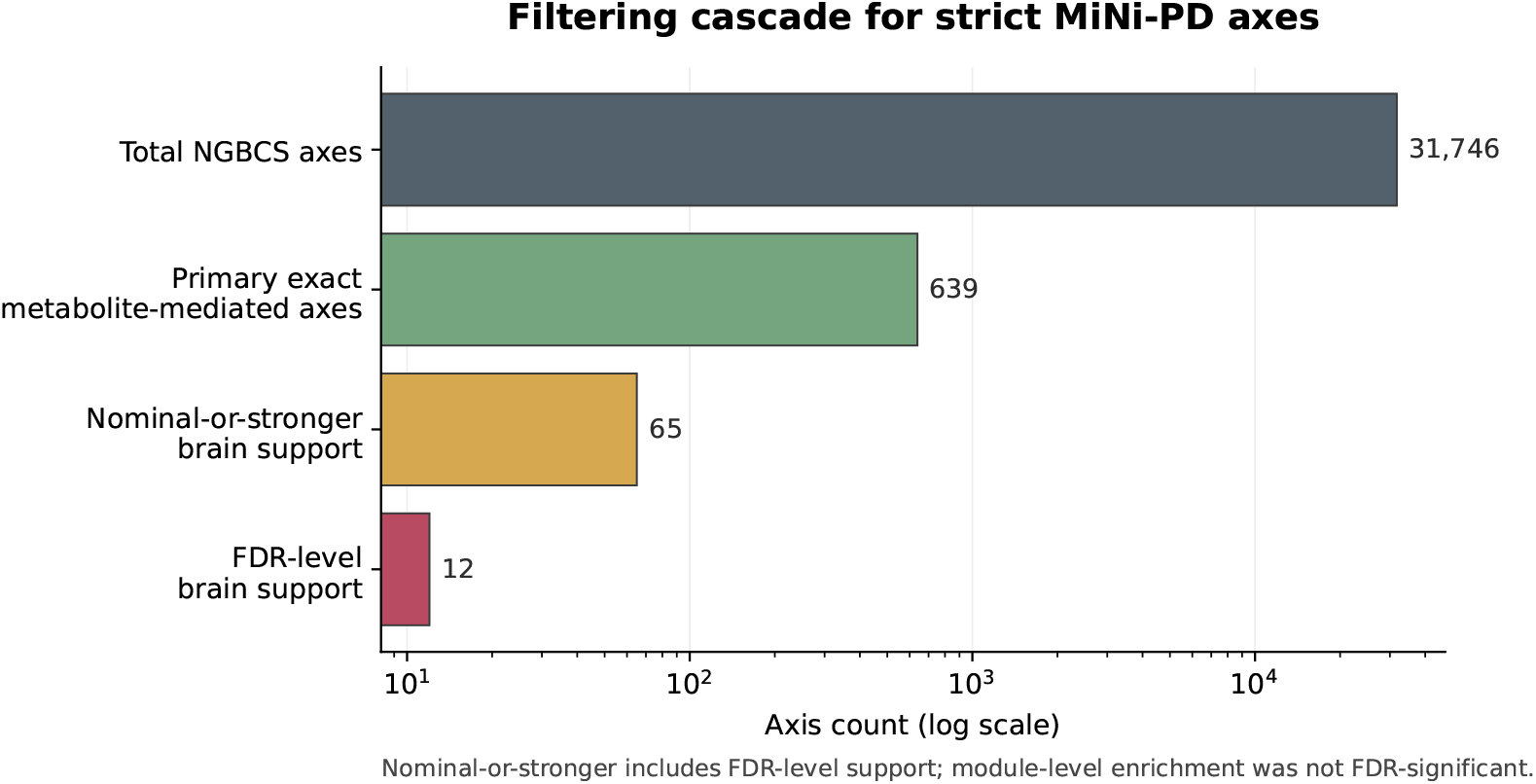
Filtering cascade for strict MiNi-PD axes. Filtering reduced 31,746 total NGBCS axes to 639 primary exact-curated metabolite-mediated axes. Among these, 12 had FDR-level brain transcriptomic support and 65 had nominal-or-stronger brain transcriptomic support. Nominal-or-stronger support includes the FDR-level subset.

**Figure 3.**
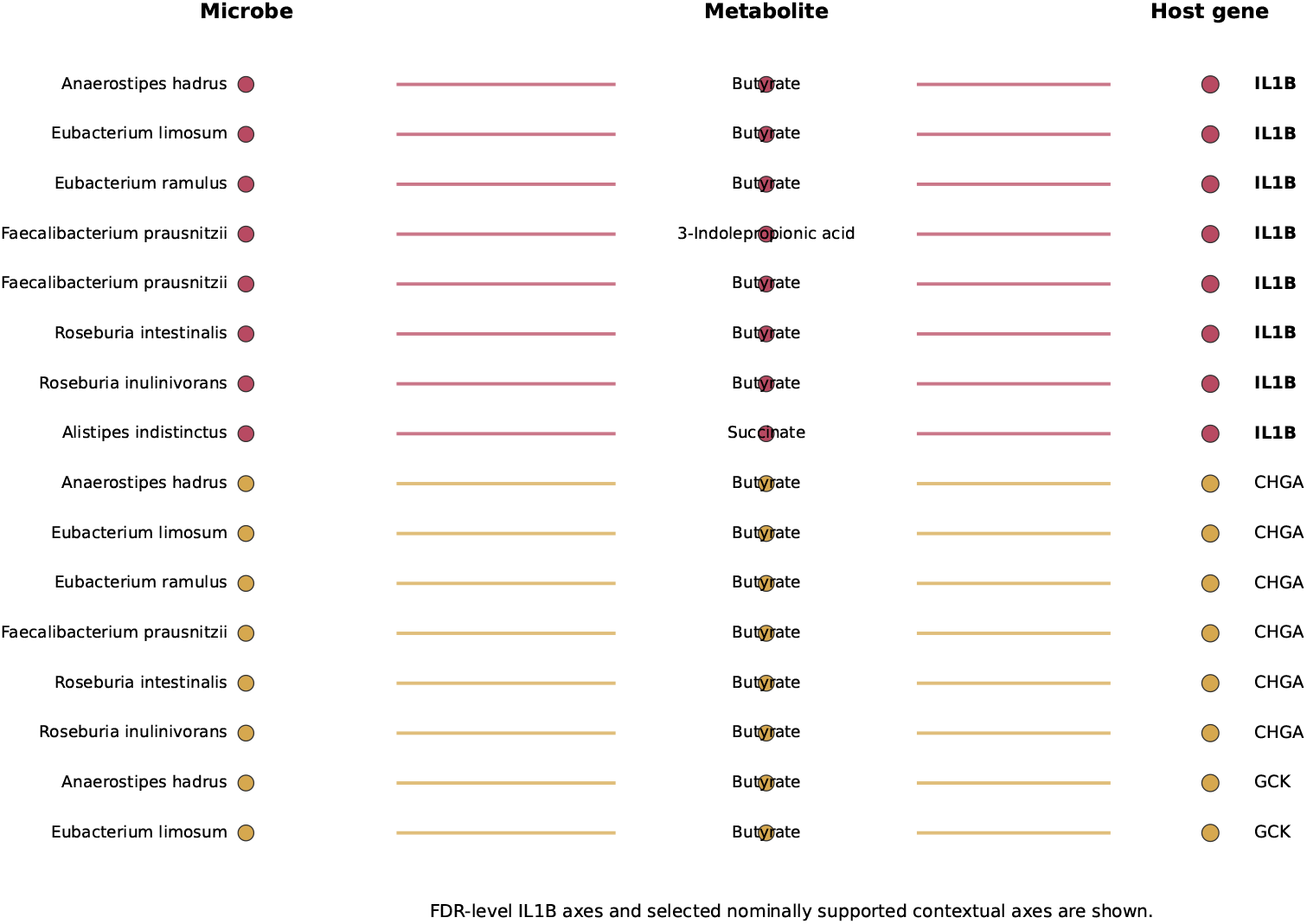
Top SCFA-linked microbe–metabolite–host-gene axes. Representative exact-curated SCFA-linked axes are shown, including FDR-level IL1B-linked axes and selected nominally supported inflammatory, metabolic, and neuroendocrine contextual axes.

**Figure 4.**
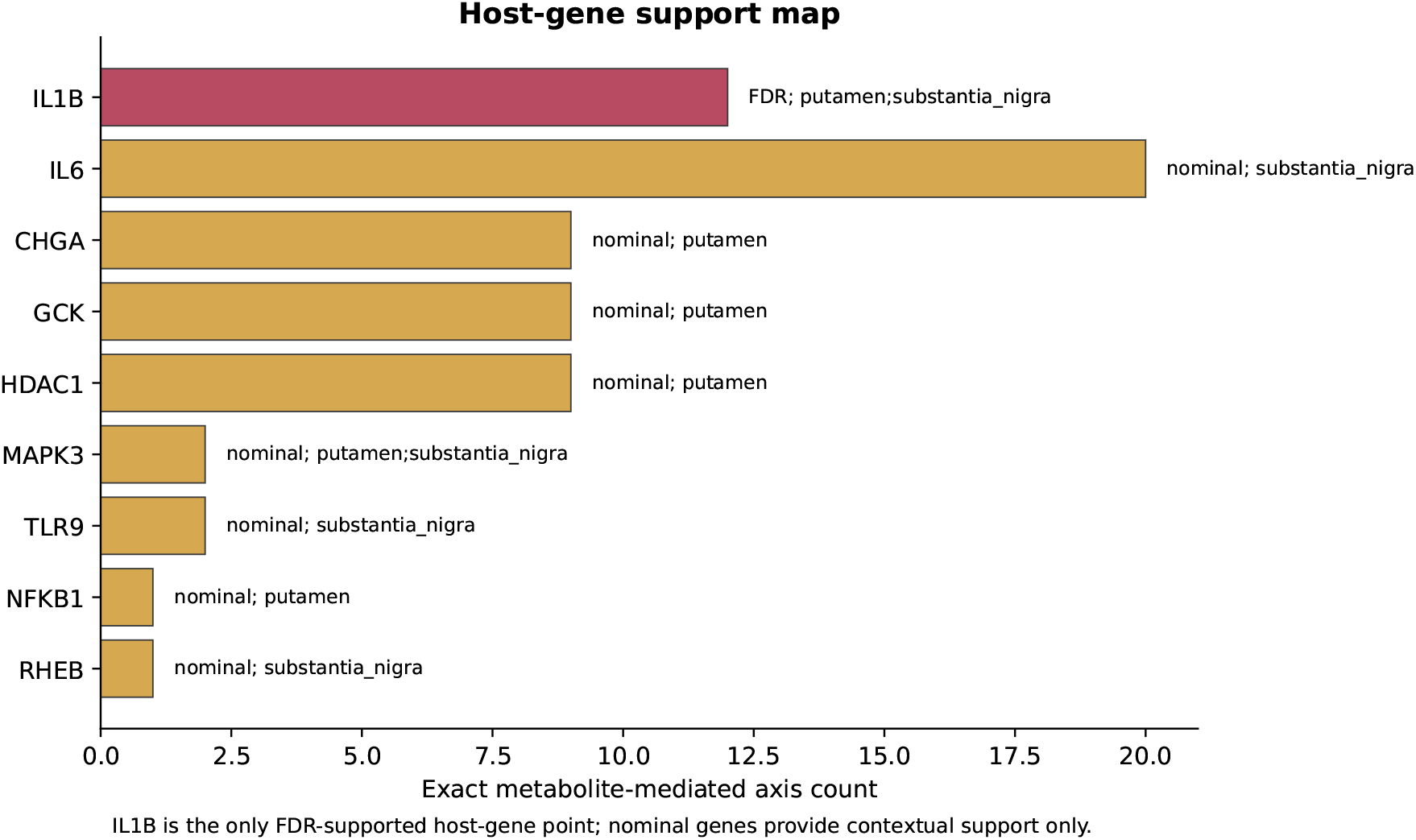
Host-gene support map centered on IL1B. Host genes are summarized by exact metabolite-mediated axis count and brain transcriptomic support tier. IL1B is the only host-gene point with FDR-level support. IL6, CHGA, GCK, HDAC1, MAPK3, TLR9, NFKB1, and RHEB provide nominal contextual support.

**Figure 5.**
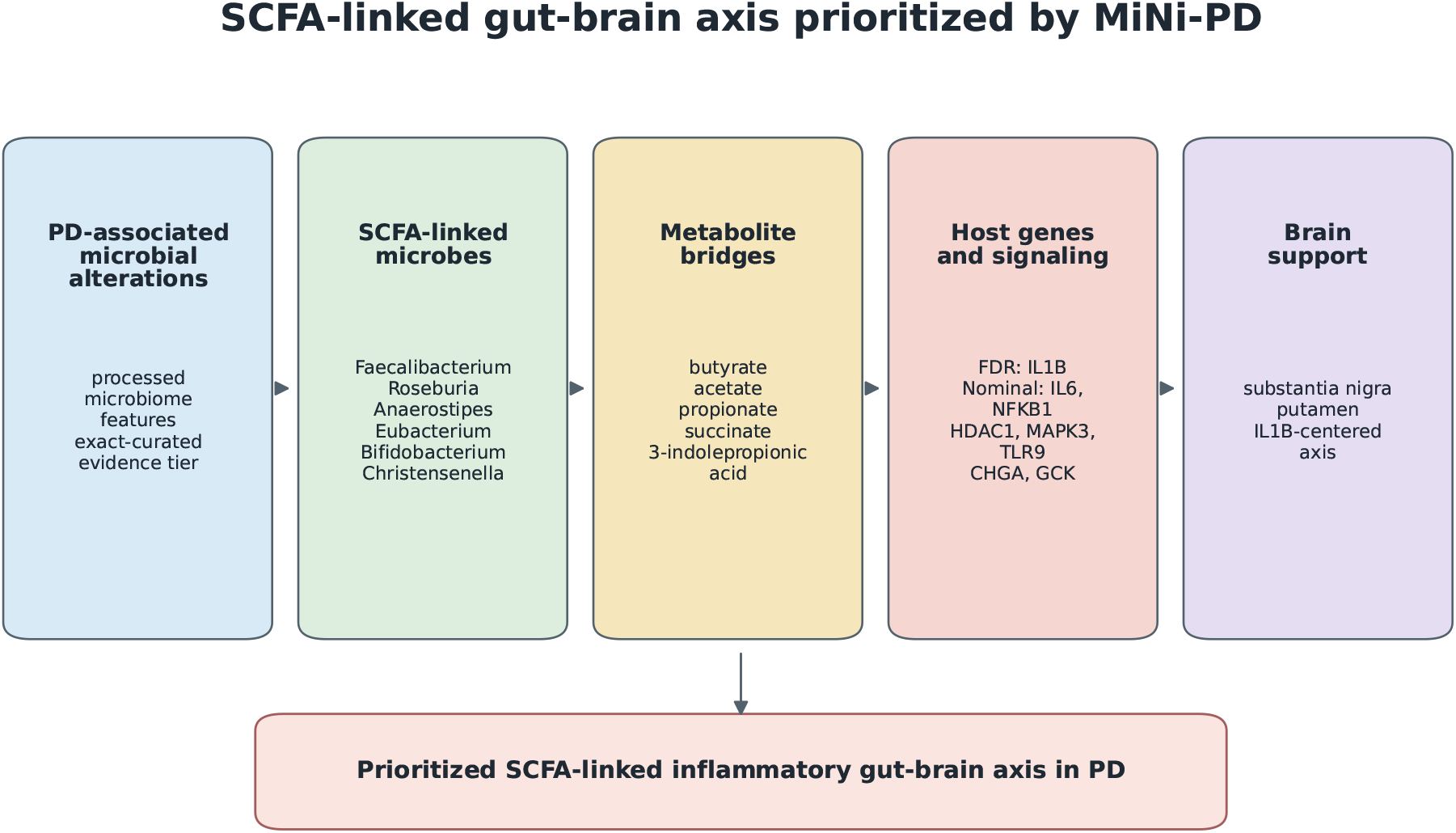
SCFA-linked IL1B-centered MiNi-PD gut-brain model. The model summarizes the central interpretation: PD-associated microbial alterations connect to SCFA-linked microbes and metabolites, host inflammatory or metabolic signaling, and brain transcriptomic support centered on IL1B. Additional nominally supported genes provide inflammatory and metabolic context.

## 15 Supplementary Material Description

**Supplementary Table S1. Genus-level axes**. This table contains genus-level curated axes generated during MiNi-PD prioritization. These rows provide broader taxonomic context and are separated from exact-curated microbial matches used for the primary evidence tier.

**Supplementary Table S2. Direct microbe–host-gene evidence without explicit metabolite mediation**. This table contains direct microbe–host-gene evidence lacking an explicit metabolite-mediated link. These rows provide supplementary biological context alongside the primary metabolite-mediated axes.

**Supplementary Table S3. Detailed metabolite-level support summary**. This table provides the detailed microbe and host-gene lists underlying the compact metabolite-level summary shown in Table 4.

**Supplementary Table S4. Detailed microbe-level support summary**. This table provides the detailed metabolite and host-gene lists underlying the compact microbe-level summary shown in Table 5.

**Source provenance and strict filtering**. Primary evidence tables were derived from processed Wallen et al. microbiome result tables, gutMGene v2.0 curated relationships, and processed GSE136666 brain transcriptomic tables. Strict filtering retained exact-curated metabolite-mediated axes and separated FDR-level support, nominal support, genus-level evidence, and direct microbe–host-gene evidence into distinct evidence tiers.

